# Repurposing Antibacterial Drugs and Clinical Molecules to Combat Vancomycin-Resistant Enterococci (VRE)

**DOI:** 10.1101/2025.03.27.645859

**Authors:** Somaia M. Abdelmegeed, Mohamed F. Mohamed, Mohamed N. Seleem

## Abstract

Vancomycin-resistant enterococci (VRE) is a growing threat to public health due to its increasing prevalence and limited treatment options. The urgent need for alternative therapeutics necessitates novel strategies such as drug repurposing to identify effective antimicrobial agents. In this study, we screened a library of 1,135 antibacterial compounds to identify candidates with activity against VRE. A total of 58 active compounds were identified, with ridinilazole and CRS3123 emerging as the most promising hits. Both compounds exhibited potent antibacterial activity in the nanomolar range against multiple *Enterococcus* species, including vancomycin-resistant and vancomycin-sensitive *E. faecium, E. faecalis, E. hirae* and *E. durans*. CRS3123 demonstrated exceptional potency, exhibiting MIC values of <0.007 μg/mL against most strains. Ridinilazole exhibited an MIC spectrum of <0.007 to 1 μg/mL across all strains, with *E. faecium* being the most sensitive (MIC: <0.007–0.25 μg/mL) and *E. faecalis* displaying a higher MIC range (0.5–1 μg/mL). Time-kill assays indicated that both compounds exhibited bacteriostatic activity comparable to linezolid. Additionally, ridinilazole and CRS3123 displayed low cytotoxicity in Vero cells and no hemolytic activity, suggesting a favorable safety profile. In vivo studies using the *C. elegans* infection model further confirmed their efficacy, with ridinilazole achieving a 60% reduction in VRE burden, CRS3123 resulting in a 25% reduction, and linezolid demonstrating a 55% reduction. These findings highlight ridinilazole and CRS3123 as promising candidates for further development for managing VRE infections.

## Introduction

Enterococci are Gram-positive, facultative anaerobic bacteria that exist naturally in the environment and are an important part of the normal gut microbiota in both humans and animals (1). Among the nearly 50 recognized species, *Enterococcus faecium* and *E. faecalis* are the most clinically significant. Enterococcal infections commonly present as urinary tract infections (UTIs), intra-abdominal infections, endocarditis, and bloodstream infections (BSIs) (2). Less frequently, they can also lead to septic arthritis, pneumonia, osteomyelitis, and meningitis. Due to their ability to exploit weakened immune systems, enterococci are considered opportunistic pathogens that are capable of causing severe and sometimes fatal infections (3).

Treating enterococcal infections pose a significant challenge due to their high level of antibiotic resistance. In particular, vancomycin-resistant enterococci (VRE) have become a major public health concern (2). According to the Centers for Disease Control and Prevention (CDC), VRE was responsible for hospitalizations of approximately 55,000 patients in the U.S. during 2017, with a mortality rate of 10% and an estimated healthcare burden of $540 million (4, 5). The increasing prevalence of VRE infections is further exacerbated by the organism’s ability to rapidly acquire resistance through genetic modifications and horizontal gene transfer. Given these concerns, the World Health Organization (WHO) has classified VRE as a high-priority multidrug-resistant pathogen, emphasizing the urgent need for novel therapeutics (6).

Currently, the treatment options for VRE infections are severely limited. Linezolid remains the only FDA-approved drug for VRE infections (7). However, its clinical use is associated with significant drawbacks, including a high mortality rate (up to 30% for bloodstream infections), limited gastrointestinal decolonization efficacy, and significant adverse effects such as bone marrow suppression and neurotoxicity (8, 9). Daptomycin, another commonly used antibiotic, is not FDA-approved for VRE infections, and its inconsistent dosing regimens raise concerns about efficacy (10). Additionally, VRE has developed resistance to multiple frontline antibiotics, including linezolid, daptomycin, quinupristin/dalfopristin, and tigecycline, further limiting treatment options (11). Given this alarming resistance trend, there is an urgent demand for new anti-VRE therapeutics (12–14).

One promising approach to address the antibiotic resistance crisis is drug repurposing, which involves discovering new therapeutic applications for already approved medications (15–17). This strategy significantly accelerates the drug discovery pipeline by leveraging compounds that have already undergone extensive safety and pharmacokinetic evaluations in humans (18). As a result, repurposed drugs bypass many of the early-stage preclinical hurdles, reducing both the time and financial investment required for drug development (19). Given the urgent need for new treatment options against multidrug-resistant pathogens like VRE, drug repurposing provides a practical and efficient pathway for discovering novel antibacterial agents (18).

Antibacterial drug repurposing focuses on screening existing compounds, including FDA-approved drugs and investigational agents, to identify those with previously unrecognized activity against resistant bacteria (19). Many of these compounds were originally developed for non-infectious diseases but have demonstrated antimicrobial properties, either through direct bacterial inhibition or by enhancing host immune responses (20). This approach is particularly valuable for combating VRE, as new antibiotics targeting enterococcal infections have been slow to emerge, and resistance to existing therapies continues to rise (18).

In this study, we conducted screening of a library of 1,135 antibacterial compounds to identify potential candidates with activity against VRE. Our goal focused on identifying compounds with potent anti-enterococcal activity, aiming to uncover alternative therapeutic options that could be rapidly transitioned into clinical use for VRE. Our findings indicate that ridinilazole and CRS3123 (previously called REP3123), two novel antibiotics currently in clinical trials for *Clostridioides difficile* infection, exhibit potent activity against VRE. These results highlight the potential of repurposed and investigational drugs in the development of effective treatments for infections caused by VRE.

## Materials and methods

### Reagents and bacterial strains

All culture media and chemical reagents were obtained from commercial suppliers: tryptic soy agar (TSA) and tryptic soy broth (TSB) were obtained from Becton, Dickinson (Cockeysville, MD). The antibacterial library (Cat. No. HY-L049) obtained from MedChemExpress (Monmouth Junction, NJ), comprising 1,135 distinct compounds with known antibacterial activity. Ridinilazole and CRS3123 were purchased from A2B Chem LLC (San Diego, CA). Linezolid (Selleck Chemicals) and vancomycin hydrochloride (Gold Biotechnology) were all purchased from commercial suppliers. Enterococcal strains were sourced from American Type Culture Collection (ATCC) (Manassas, VA, USA) and the Biodefense and Emerging Infections Research Resources Repository (BEI Resources) (Manassas, VA, USA).

### Screening assay for enterococci

A single round of screening was performed on an antibacterial compound library using *Enterococcus faecium* NR-31909 strain. The broth microdilution was conducted in accordance with the Clinical and Laboratory Standards Institute (CLSI) guidelines (16, 20). In brief, *Enterococcus faecium* NR-31909 was grown on tryptic soy agar (TSA) followed by incubation at 37°C for 24 hours. A bacterial suspension was adjusted to 0.5 McFarland standard and suspended in tryptic soy broth (TSB) to obtain 5 × 10⁵ CFU/mL concentration, then dispensed into 96-well plates with addition of a final concentration of 4 µM of the test compounds. Then the plates were kept in the incubator at 37°C for 24 hours. Negative control, consisting of dimethyl sulfoxide (DMSO), was included in the experiment, while vancomycin and linezolid were used as reference antibiotics. The percentage of bacterial growth inhibition was determined for each compound, and compounds that exhibited at least 80% inhibition were classified as “hits.” Active hits were confirmed, and the actual minimum inhibitory concentration (MIC) readings were measured by the broth microdilution technique (16, 20). The MIC was identified as the lowest concentration at which bacterial growth was completely inhibited. Data visualization, including the percentage of bacterial inhibition, was generated using GraphPad Prism version 10.

### Antibacterial activity of the potent hits against various clinical enterococci strains

Two potent compounds (ridinilazole and CRS3123) were selected based on nanomolar activity against VRE for additional analysis. These two compounds were obtained commercially and tested against a group of 15 clinical isolates of enterococci, comparing their efficacy to the control antibiotics, vancomycin and linezolid by the broth microdilution technique as mentioned above (16, 20).

### Time-kill assay

The time-kill assay was performed as previously published (21). In TSB, *E. faecium* NR-31909 was cultured overnight. Then suspended in fresh TSB and kept at 37°C with aerobic incubation until the cells were in the logarithmic phase, as indicated by an optical density (OD₆₀₀) of 0.2. The bacterial cultures were further diluted to adjust to a final concentration of 5 × 10⁵ CFU/mL in TSB. Linezolid, ridinilazole and CRS3123, were introduced at concentrations of 5× and 10× MIC. A DMSO-treated culture served as an untreated control. The cultures were aerobically maintained at 37°C in shaking incubator. At specific sampling times, aliquots were collected, resuspended in phosphate-buffered saline (PBS) and distributed to three replicates on tryptic soy agar (TSA). After a 24-hour incubation period at 37°C, the number of colony-forming units (CFUs) was evaluated.

### Cytotoxicity assessment in Vero cells

The potential cytotoxic effect of the test compounds was investigated using Vero cells (monkey kidney epithelial cells) following established protocols(22). Briefly, Vero cells were plated into 96-well plates supplemented with Dulbecco’s Modified Eagle Medium (DMEM) containing 10% fetal bovine serum (FBS). Then the cells placed in a humidified 37°C environment for incubation with 5% CO₂ for 24 hours to allow proper adherence and growth. Following the incubation period, we replaced the culture medium with fresh DMEM containing serial dilutions of the test compounds, while the solvent alone used as a negative control. The treated cells were further incubated for another 24 hours. To assess cell viability, the MTS/PMS reagent was added to each well four hours and a microplate reader (Synergy H1, BioTek, USA) was used to measure absorbance at 490 nm. All experiments were conducted in three biological replicates.

### Hemolysis assay

The hemolytic activity of the compounds was evaluated as previously described (23, 24). Fresh human red blood cells (RBCs) (Innovative Research, Novi, MI) were centrifuged at 2,000 rpm for 5 minutes to form a pellet. The RBC pellet was rinsed with PBS three times. An 8% (v/v) suspension of red blood cells (RBCs) was suspended in PBS and 50 µL of this suspension was added to each well of a 96-well plate. 50 µL of the test compounds at varying concentrations, prepared in PBS, were added to achieve RBC concentration of 4% (v/v) in each well. PBS alone served as the negative control, while 0.1% Triton X-100 was used as the positive control. The plate was kept at 37°C incubator for 1 hour, followed by centrifugation at 1,000 rpm for 5 minutes at 4°C to separate the intact RBCs from the supernatant. Carefully, 75 µL of the resultant supernatants were moved to a new sterile 96-well plate. Hemolysis was quantified using a microplate reader (Synergy H1, BioTek, USA) by measuring the absorbance at 405 nm. The percentage of hemolysis was calculated relative to the 100% hemolysis control (0.1% Triton X-100). Experiments were conducted in three biological replicates.

### *In vivo* efficacy of ridinilazole and CRS3123 in a *Caenorhabditis elegans* (*C. elegans*) infection model

The temperature-sensitive sterile mutant strain *C. elegans* AU37 [sek-1(km4); glp-4(bn2) I] was applied to evaluate the in vivo efficacy of drugs, as previously described (23, 25–28). In summary, adult worms were cultured on NGM agar plates inoculated with a lawn of *E. coli* OP50at 15°C for 5 days to allow egg-laying. The eggs were bleached to harvest them and incubated at room temperature for 24 hours with gentle agitation. to facilitate hatching. Newly hatched larvae were placed to fresh NGM plates containing *E. coli* OP50 and maintained at room temperature until they matured to the adult stage. Once the worms matured, they were gathered and rainsed three times with PBS to eliminate residual *E. coli* before being infected with *E. faecium* NR-31909. Following infection, the worms were washed three times with PBS. Approximately 30 worms per sample were treated with 10x MIC of the test compounds for 24 hr. To determine the bacterial load, worms were rinsed three times using PBS and then lysed by adding to each tube 200 mg of 1.0-mm silicon carbide particles (Biospec Products, Bartlesville, OK), followed by vortexing for five minutes. The resulting bacterial solution was plated onto TSA plates supplemented with ampicillin and vancomycin to select VRE, and incubated at 37°C for 24 hours, and bacterial colonies were subsequently counted.

## Results

### Screening of the antibacterial library against enterococci

To identify potential novel anti-enterococci drugs, 1135 antibacterial compounds were screened for their activity using vancomycin resistant *E. faecium* NR-31909. The initial screening was performed at a concentration of 4 μM. Of these, 58 compounds demonstrated anti-enterococci activity by effectively inhibiting the enterococci growth at the tested concentration of 4 μM. (Fig. 1). The hits were further screened to detect the actual MIC using the broth microdilution method. The MIC measurements of the resulting hits was ranging from <0.031 to 4 μM (Table 1). The two most promising compounds, ridinilazole and CRS3123, which exhibit anti-enterococci activity at nanomolar range (<0.031 μM) were selected for further investigation.

**Figure 1:**
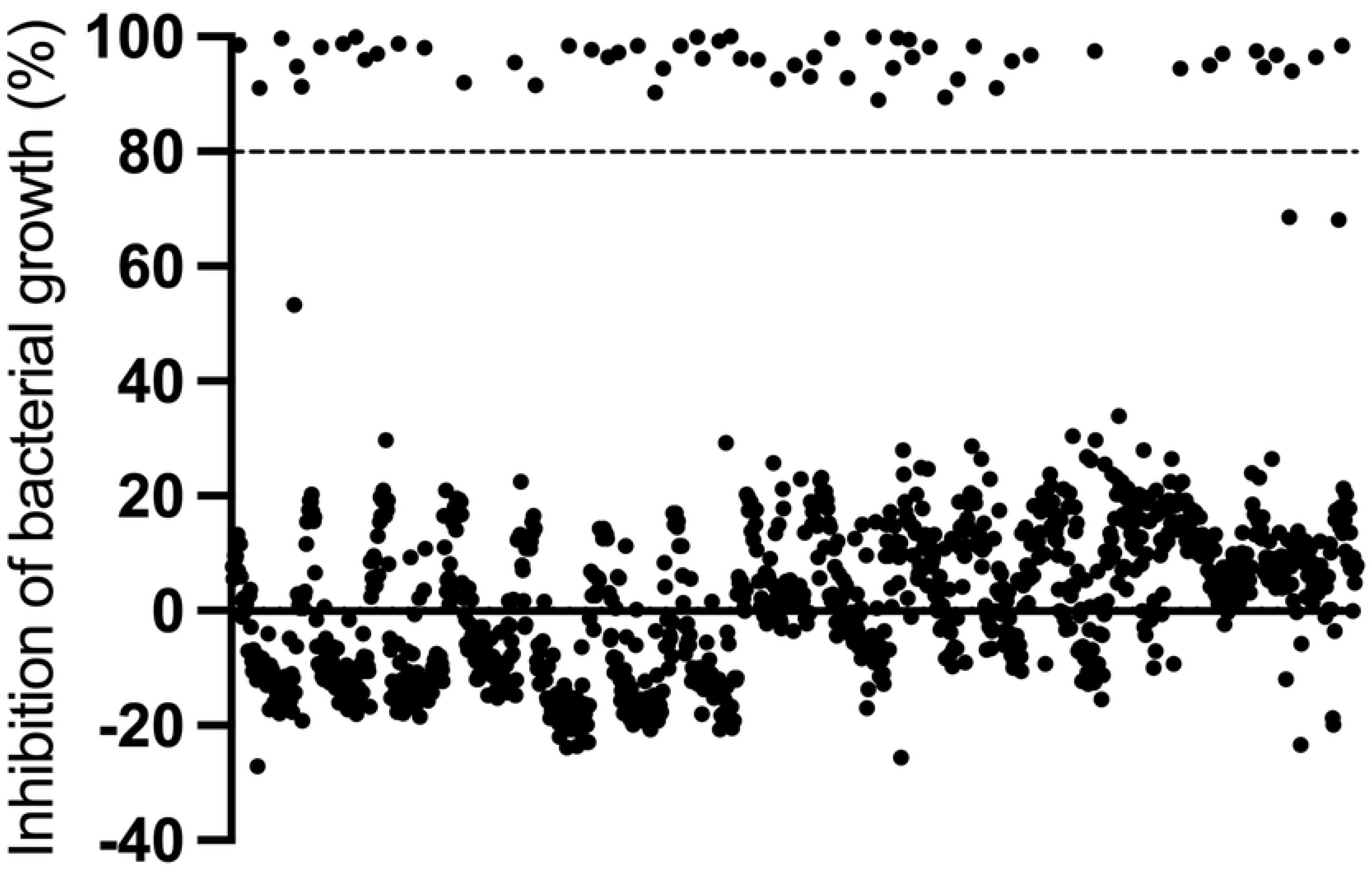
MCE antibacterial library screening of 1135 compounds at 4 μM against *E. faecium* NR-31909. Compounds with a greater than 80% inhibition of bacterial growth were identified as hits while hits below 80% inhibition were excluded due to lack of activity. 58 total hits were identified.

**Table 1.** Description and MICs (μM) of potent antibacterial library hits against *E. faecium* NR-31909.

| Compound ID | MIC | Description | Clinical information |
| --- | --- | --- | --- |
| Walrycin B | 4 | Antibiotic; Bacterial | No Development Reported |
| Gliotoxin | 0.25 | Antibiotic; Apoptosis; Bacterial; Fungal; NF-κB; PKA | No Development Reported |
| Tiamulin (fumarate) | 1 | Antibiotic; Bacterial | No Development Reported |
| PA3552-IN-1 | 4 | Bacterial | No Development Reported |
| BRITE-338733 | 4 | Bacterial | No Development Reported |
| Thonzonium (bromide) | 4 | Bacterial; Proton Pump | Launched |
| Anacardic Acid | 4 | Bacterial; Epigenetic Reader Domain; Histone Acetyltransferase | No Development Reported |
| Cetalkonium (chloride) | 4 | Bacterial | Launched |
| Cetylpyridinium (chloride) | 4 | Bacterial; HBV | Launched |
| Ethoxzolamide | 2 | Bacterial; Carbonic Anhydrase | Launched |
| Mupirocin | 2 | Antibiotic; Bacterial | Launched |
| alpha-Mangostin | 4 | Apoptosis; Bacterial; Fungal; Reactive Oxygen Species; Virus Protease | No Development Reported |
| Ridinilazole | <0.031 | Bacterial | Phase 3 |
| Acetazolamide | 1 | Autophagy; Bacterial; Carbonic Anhydrase | Launched |
| Brilacidin | 2 | Antibiotic; Bacterial | Phase 2 |
| Fusidic acid | 2 | Antibiotic; Bacterial | Launched |
| Maduramicin (ammonium) | 2 | Antibiotic; Apoptosis; Bacterial | No Development Reported |
| Novobiocin (sodium) | 2 | Antibiotic; Apoptosis; Bacterial; DNA/RNA Synthesis; HSP; Orthopoxvirus | Launched |
| Nanchangmycin | 4 | Antibiotic; Bacterial | No Development Reported |
| Nosiheptide | 0.5 | Antibiotic; Bacterial | No Development Reported |
| Nigericin (sodium salt) | 0.5 | Antibiotic; Bacterial; NOD-like Receptor (NLR); Potassium Channel | No Development Reported |
| Pristinamycin | 2 | Antibiotic; Bacterial | Phase 4 |
| Tigecycline (tetramesylate) | 1 | Antibiotic; Autophagy; Bacterial | Launched |
| 1-Monomyristin | 4 | Autophagy; Bacterial; Endogenous Metabolite; FAAH; Fungal | No Development Reported |
| Antibacterial agent 18 | 2 | Bacterial | No Development Reported |
| Auranofin | 2 | Bacterial; SARS-CoV | Launched |
| Cadazolid | 1 | Antibiotic; Bacterial | Phase 3 |
| Radezolid | 2 | Antibiotic; Bacterial | Phase 2 |
| Retapamulin | 0.25 | Antibiotic; Bacterial | Launched |
| Pleuromutilin | 4 | Antibiotic; Bacterial | No Development Reported |
| Cetylpyridinium (chloride monohydrate) | 4 | Bacterial | Launched |
| Contezolid | 4 | Antibiotic; Bacterial; Monoamine Oxidase | Launched |
| SPR719 | 0.125 | Bacterial; Topoisomerase | No Development Reported |
| Mupirocin (calcium hydrate) | 1 | Antibiotic; Bacterial | Launched |
| NH125 | 4 | Autophagy; Bacterial; CaMK; Fungal; Virus Protease | No Development Reported |
| Tridecanoic acid | 4 | Bacterial; Endogenous Metabolite | Phase 2 |
| Dianemycin | 2 | Antibiotic; Bacterial | No Development Reported |
| Bithionol | 2 | Bacterial; Parasite | Launched |
| Valnemulin (hydrochloride) | 0.125 | Antibiotic; Bacterial | No Development Reported |
| Calcimycin | 4 | Antibiotic; Apoptosis; Autophagy; Bacterial; Fungal | Phase 4 |
| Omadacycline (hydrochloride) | 4 | Antibiotic; Bacterial | Launched |
| Virginiamycin M1 | 4 | Antibiotic; Bacterial | No Development Reported |
| CRS3123 (dihydrochloride) | <0.031 | Antibiotic; Bacterial | Phase2 |
| Gramicidin | <0.031 | Antibiotic; Bacterial | Launched |
| Chromomycin A3 | 4 | Antibiotic; Apoptosis; Bacterial; Fungal | No Development Reported |
| Oritavancin (diphosphate) | 0.5 | Antibiotic; Bacterial | Launched |
| Trimethyloctadecylammonium (bromide) | 2 | Dynamin | No Development Reported |
| Floxuridine | 1 | Apoptosis; Bacterial; CMV; DNA/RNA Synthesis; HSV; Nucleoside Antimetabolite/Analog | Launched |
| Lasalocid | 1 | Antibiotic; Autophagy; Bacterial; Parasite | No Development Reported |
| MCB-3681 | 4 | Bacterial | No Development Reported |
| Thiostrepton | 4 | Antibiotic; Bacterial | No Development Reported |
| Clofoctol | 4 | Antibiotic; Bacterial; SARS-CoV | Launched |
| Acetazolamide (sodium) | 1 | Autophagy; Bacterial; Carbonic Anhydrase | Launched |
| Lasalocid (sodium) | 1 | Antibiotic; Autophagy; Bacterial; Parasite | No Development Reported |
| Reutericyclin | 4 | Antibiotic; Bacterial | No Development Reported |
| Fidaxomicin | 4 | Antibiotic; Apoptosis; Bacterial; DNA/RNA Synthesis | Launched |
| Decamethoxine | 4 | Bacterial; Fungal | No Development Reported |
| Octenidine (dihydrochloride) | 1 | Bacterial | Launched |

### Antibacterial activity of the potent two hits against various clinical enterococci strains

The antibacterial effect of the potent two hits, ridinilazole and CRS3123 was investigated further by testing against 15 enterococci clinical isolates including vancomycin resistant and sensitive *E. faecium*, *E. faecalis, E. durans,* and *E. hirae.* (Table 2). The MIC of CRS3123 were <0.007 μg/mL against all strains except *E. faecalis* HM-335 (MIC=0.031 μg/mL). The MIC of ridinilazole ranged from <0.007 to 1 μg/mL against all strains, with *E. faecium* being the most sensitive (MIC range: <0.007–0.25 μg/mL), while *E. faecalis* exhibited a higher MIC range (0.5–1 μg/mL). The MIC of linezolid was ranging from 0.5-2 μg/mL, while most strains showed resistance to vancomycin (Table 2).

**Table 2.** MIC values (μg/mL) of the most potent hits (CRS3123 and ridinilazole) against clinical isolates of enterococci.

| Strain | CRS3123 | Ridinilazole | Linezolid | Vancomycin |
| --- | --- | --- | --- | --- |
| *Enterococcus faecium* NR-31909 | <0.007 | <0.007 | 0.5 | >128 |
| *Enterococcus faecium* -- ERV165, HM-970. | <0.007 | 0.015 | 0.5 | >128 |
| *Enterococcus faecium* NR-31912. | <0.007 | 0.063 | 0.5 | >128 |
| *Enterococcus faecium* -- E2620 NR-31954 | <0.007 | 0.031 | 2 | 0.5 |
| *Enterococcus faecium* -- E1604. NR-31935 | <0.007 | 0.031 | 0.5 | 0.5 |
| *Enterococcus faecium* -- E417 HM-965 | <0.007 | 0.015 | 2 | >128 |
| *Enterococcus faecium* – 503 HM-952. | <0.007 | 0.031 | 0.5 | >128 |
| *Enterococcus faecium* -- UAA945 NR-32094 | <0.007 | 0.25 | 0.5 | >128 |
| *Enterococcus faecalis* -- B3336. NR-31887 | <0.007 | 1 | 0.5 | >128 |
| *Enterococcus faecalis* -- R712. HM-335. | 0.031 | 1 | 0.5 | 128 |
| *Enterococcus faecalis* NR31971 2 | <0.007 | 0.5 | 0.5 | 32 |
| *Enterococcus faecalis* NR31970. | <0.007 | 1 | 0.5 | 0.5 |
| *Enterococcus faecalis* NR31972. | <0.007 | 0.5 | 0.5 | >128 |
| *Enterococcus hirae* ATCC 10541 | <0.007 | 0.125 | 2 | 0.5 |
| *Enterococcus durans* ATCC 11576 | <0.007 | <0.007 | 0.5 | 1 |

### Time-kill kinetics against enterococci

To determine whether the ridinilazole and CRS3123 have bacteriostatic or bactericidal activity against enterococci, a time-kill assay was conducted against *E. faecium* NR-31909. As shown in Fig 2, both ridinilazole and CRS3123 at 5× and 10× MIC exhibited bacteriostatic activity similar to linezolid.

**Figure 2:**
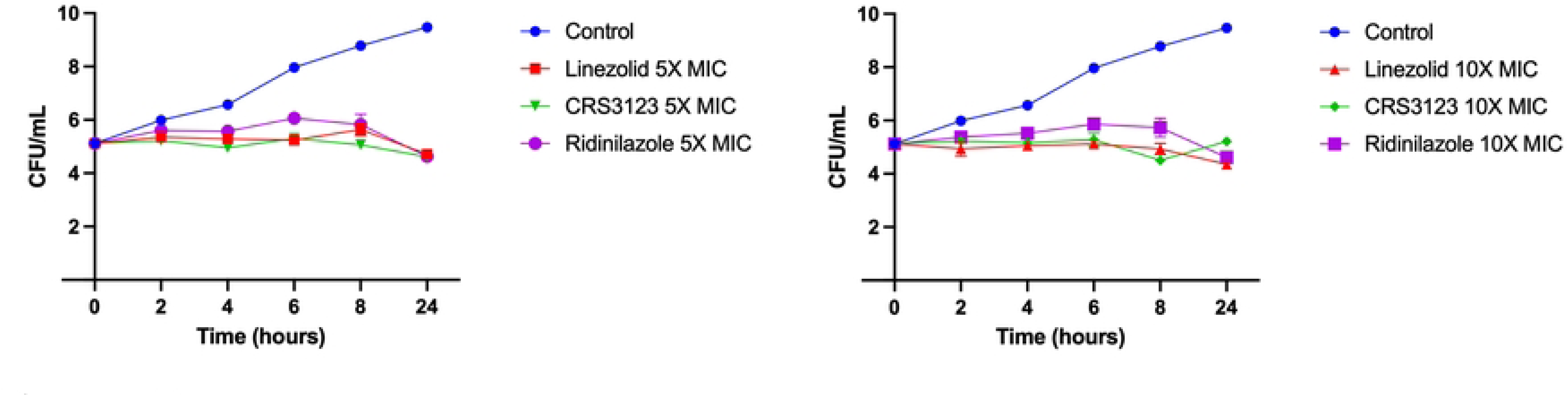
Time kill assay of ridinilazole, CRS3123 and linezolid at 5× and 10× MIC, against *E. faecium* NR-31909. Samples treated with DMSO were used as negative control. The results are given as means ± SD (n = 3; data without error bars indicate that the SD is too small to be seen).

### Cytotoxicity assessment in Vero Cells

The cytotoxicity of CRS3123 and ridinilazole was evaluated in Vero cells to assess potential toxic effects. CRS3123 demonstrated no toxicity up to 16 µg/mL but exhibited 20% cytotoxicity at 32 µg/mL, indicating mild toxicity at this concentration (Fig. 3). In contrast, ridinilazole showed no cytotoxicity up to the highest concentration tested, 128 µg/mL. (Fig. 3). Based on the IC50-to-MIC50 comparison, CRS3123 has a therapeutic index of >4571, while ridinilazole has a therapeutic index of >256, highlighting the strong safety profile and high selectivity of both compounds for bacterial cells over mammalian cells.

**Figure 3:**
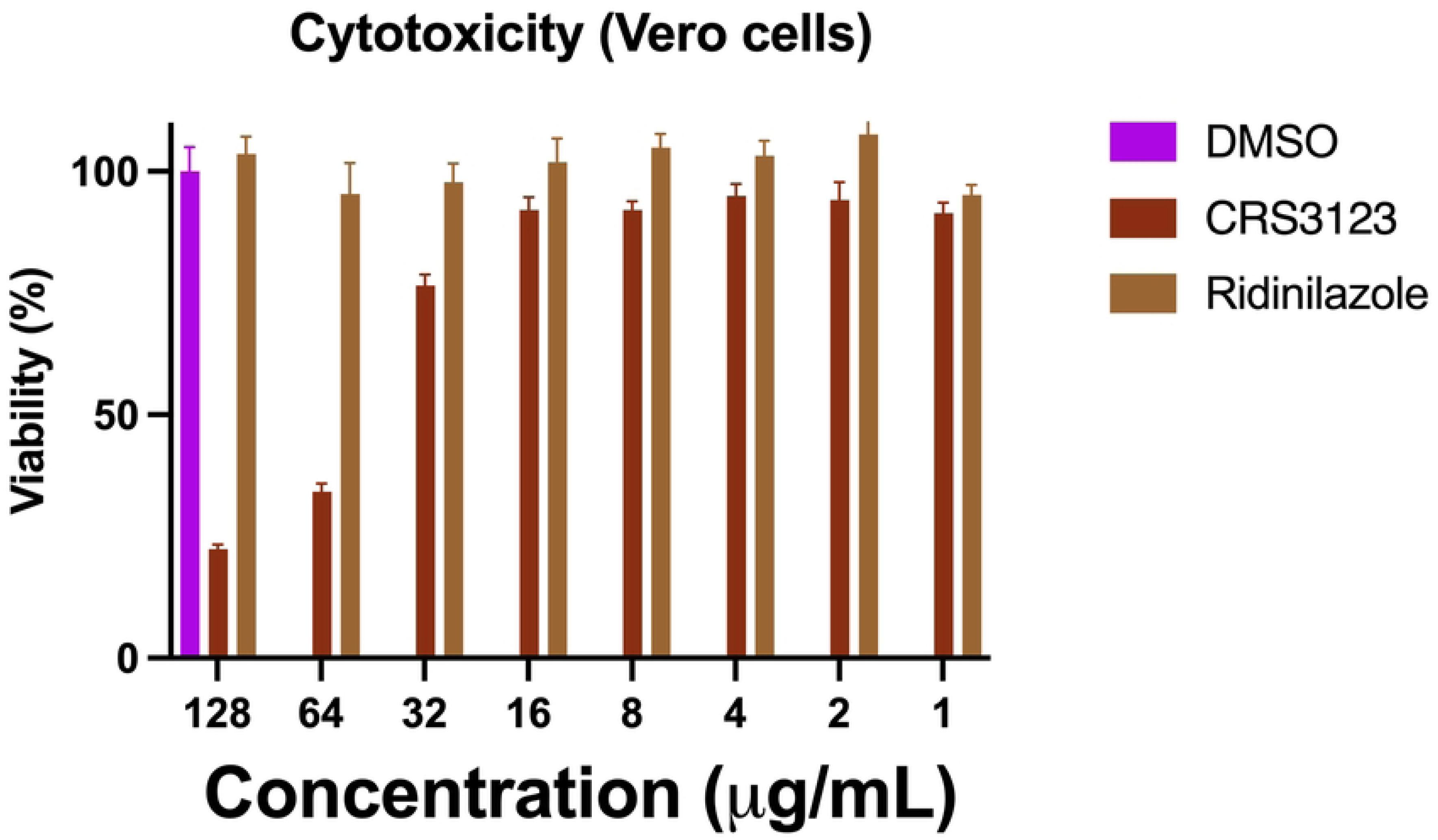
Cytotoxicity assay showing the percent mean absorbance at 490 nm after incubating Vero cells with CRES3123 and ridinilazole at different concentrations for 24 hr. Cell viability was measured by MTS assay. Results are expressed as means from three measurements ± standard deviations.

### Hemolysis assay

The hemolysis assay was conducted to assess the potential hemolytic activity of CRS3123 and ridinilazole on human red blood cells (RBCs). Both compounds exhibited no hemolytic activity up to the highest tested concentration of 256 µg/mL (Fig. 4). These results indicate that CRS3123 and ridinilazole have a strong safety profile and demonstrate high selectivity for bacterial cells over mammalian cells.

**Figure 4:**
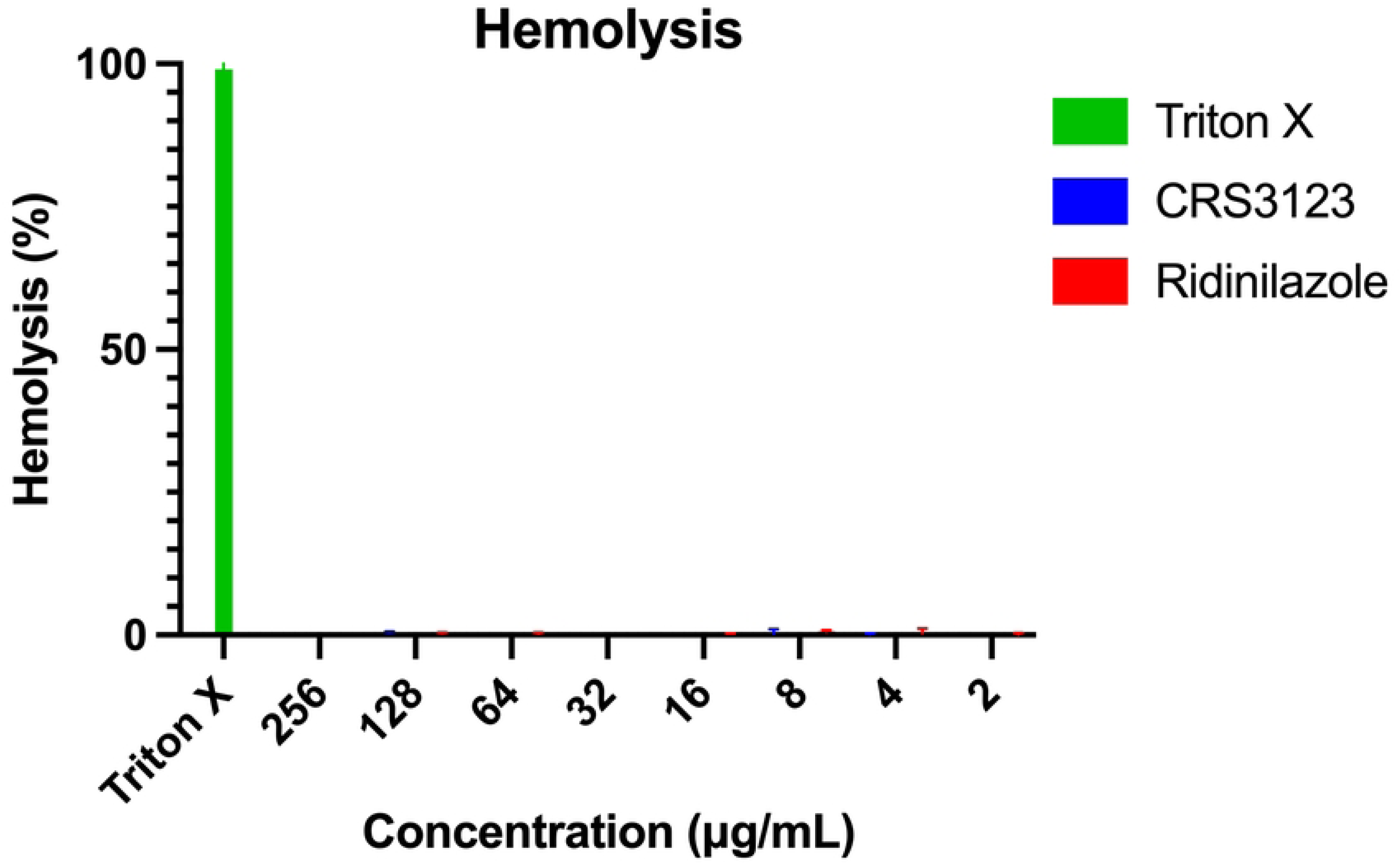
Hemolysis assay of CRS3123 and ridinilazole on human RBCs. The release of hemoglobin in the supernatant of human erythrocytes after treatment with increasing amounts of the two compounds was measured at 405 nm. Data collected after 1 h of incubation are presented. 0.1% of Triton X-100 served as positive control.

### *In vivo* efficacy of ridinilazole and CRS3123 in a *C. elegans* infection model

Building on the promising *in vitro* results, we sought to evaluate the *in vivo* efficacy of ridinilazole and CRS3123 using a *C. elegans* infection model (23, 25–28). *C. elegans* were infected with *E. faecium* NR-31909 and subsequently treated with ridinilazole, CRS3123 and linezolid at 10× MIC. After 24 hours, worms were lysed, and bacterial counts were determined. As shown in Fig. 5, ridinilazole showed 60% bacterial reduction of VRE (*p* < 0.0001), while CRS3123 resulted in 25% bacterial reduction (*p* < 0.01). The drug of choice, linezolid showed 55% bacterial reduction (*p* < 0.0001). These data demonstrating a profound *in vivo* activity of ridinilazole and CRS3123.

**Figure 5:**
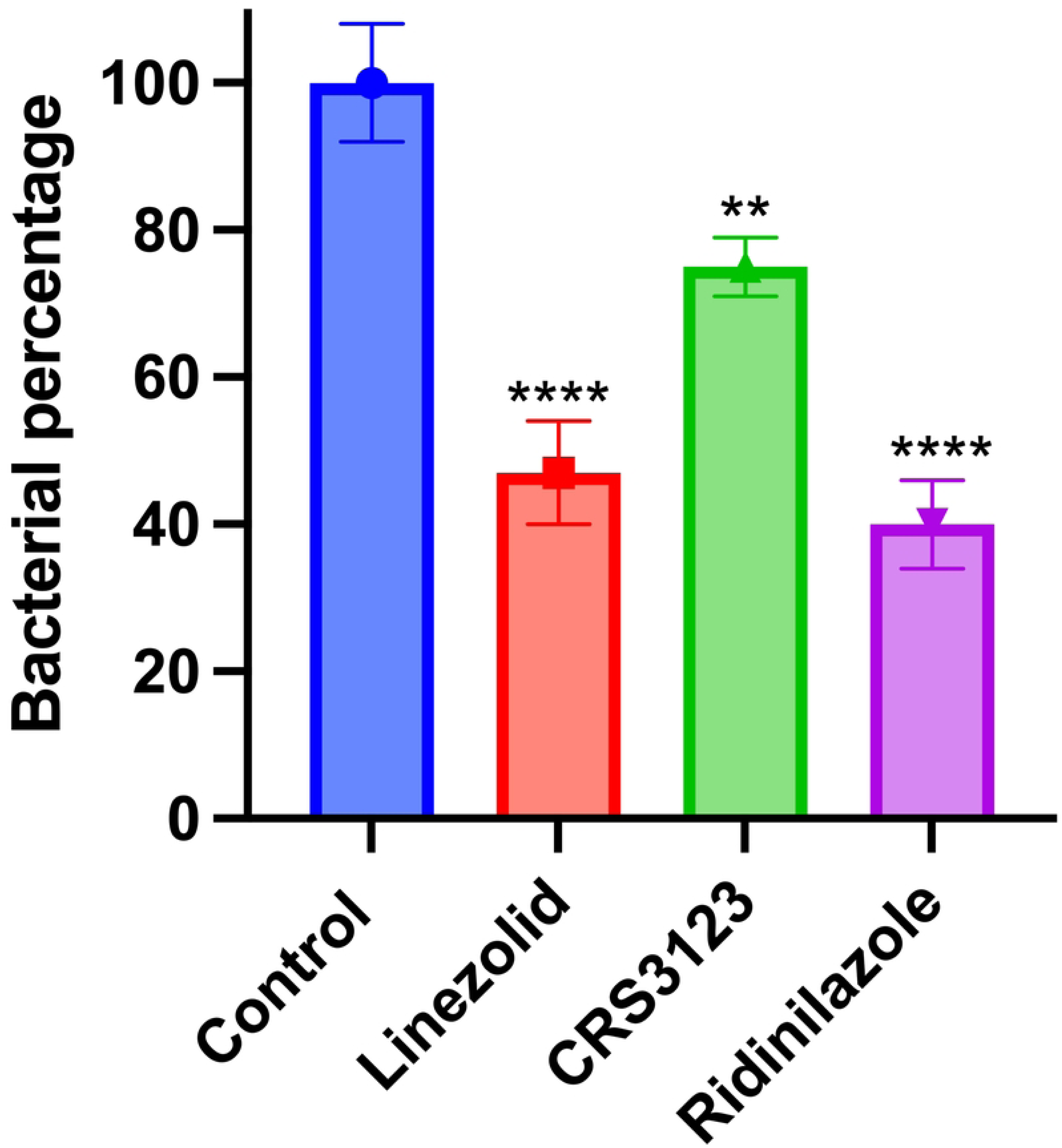
*In vivo* efficacy of ridinilazole and CRS3123 in a *Caenorhabditis elegans* model of bacterial infection. *C. elegans* were infected with VRE*, E. faecium* NR-31909. After infection, worms were treated with ridinilazole and CRS3123 or linezolid at 10× MIC. After 24 hours, worms were lysed, and bacteria were plated and CFU were counted after 24 hr. Results are expressed as means from three biological replicates ± standard deviation. Statistical analyses were determined by one-way ANOVA with post hoc testing (\*\*\**p*<0.005), (\*\*\*\**p* < 0.0001).

## Discussion

VRE infections remain a significant clinical challenge due to the increasing prevalence of resistance to last-line antibiotics, including linezolid and daptomycin (2). The need for novel therapeutics with distinct mechanisms of action is crucial to combat these infections effectively (13). CRS3123 and ridinilazole are two promising candidates with novel mechanisms, initially developed for *C. difficile* infection (CDI) (29, 30), but their activity against VRE has not been extensively characterized. Our study provides strong evidence of their potent antimicrobial effects against VRE, highlighting their potential as alternative treatment options.

Our findings demonstrate that CRS3123 exhibited strong activity against VRE, with MIC values ranged from 0.031 to 0.007 µg/mL. Compared to standard VRE treatments such as linezolid (MIC range: 0.5-2 µg/mL), CRS3123 showed higher potency. Notably, CRS3123 exhibited bacteriostatic effects at high concentration similar to linezolid. The bacteriostatic activity of CRS3123 is attributed to its selective inhibition of methionyl-tRNA synthetase, which is a crucial enzyme for bacterial protein synthesis (30). By preventing the incorporation of methionine into nascent proteins, CRS3123 disrupts bacterial translation, halting growth without directly causing cell lysis (31). This unique mechanism differentiates CRS3123 from ribosome-targeting antibiotics like linezolid, potentially reducing the risk of cross-resistance with existing protein synthesis inhibitors (31).

Similarly, ridinilazole displayed strong activity against VRE, with MIC ranged from 0.007-1µg/mL. The compound exhibited bacteriostatic effects at high concentrations. Ridinilazole disrupts bacterial cell division, mechanisms that have been well-characterized in *C. difficile* but appear to extend to enterococcal species. Ridinilazole’s interference with DNA replication and septum formation leads to arrested bacterial growth rather than bacterial killing (32). This bacteriostatic nature suggests that while ridinilazole is highly effective at controlling VRE proliferation, it may require host immune clearance or combination therapy for complete bacterial eradication (33).

The cytotoxicity and hemolysis data indicate that both CRS3123 and ridinilazole exhibit strong safety profiles with high selectivity for bacterial cells over mammalian cells. While CRS3123 demonstrated mild cytotoxicity at 32 µg/mL, its therapeutic index remains high (>4571), suggesting a favorable margin for clinical use. Ridinilazole, with no observed cytotoxicity up to 128 µg/mL and a therapeutic index >256, further supports its safety. Additionally, the absence of hemolytic activity for both compounds at concentrations up to 256 µg/mL reinforces their potential as selective and well-tolerated antibacterial agents.

Furthermore, in vivo studies using the *C. elegans* infection model confirmed the efficacy of these compounds in reducing bacterial burden. Ridinilazole demonstrated the highest reduction, decreasing VRE levels by 60%, followed by linezolid, the standard treatment, which achieved a 55% reduction. CRS3123 exhibited a more modest reduction of 25%, suggesting potential but lower efficacy in this model. These findings highlight the differential activity of these compounds in an invertebrate host system, underscoring the need for further validation in mammalian models. Future studies in murine infection models are warranted to assess pharmacokinetics, optimal dosing, and therapeutic potential against VRE in a more physiologically relevant context. Such investigations will be critical to determining the translational potential of CRS3123 and ridinilazole for clinical application.

An important factor influencing the clinical applicability of these compounds is their pharmacokinetic profile. CRS3123 achieves high concentrations in the gastrointestinal tract with minimal systemic absorption (34), which is advantageous for treating gastrointestinal infections but may limit efficacy in systemic VRE infections. This property suggests that CRS3123 may be particularly effective in gut decolonization infections (34), reducing VRE colonization in high-risk patients and preventing bloodstream infections. Further modifications to enhance systemic bioavailability may expand its therapeutic potential beyond localized infections.

Similarly, ridinilazole has been associated with minimal disruption of gut microbiota, making it an attractive candidate for treating enteric VRE infections while reducing the risk of secondary infections, such as *C. difficile* overgrowth (35). However, its systemic penetration remains an area of further investigation, as enhanced distribution into deep tissues and the bloodstream would be necessary for treating invasive VRE infections. Future pharmacokinetic studies and formulation improvements may optimize its systemic efficacy.

Another key aspect of CRS3123 and ridinilazole’s potential clinical application is their low propensity for resistance development (33, 36). Given the increasing prevalence of resistance to conventional VRE treatments, assessing the likelihood of resistance emergence is critical (34, 35). This characteristic may position CRS3123 and Ridinilazole as valuable options for combination therapy, potentially enhancing the efficacy of existing antibiotics like daptomycin or linezolid while reducing the risk of resistance selection.

In conclusion, these findings highlight ridinilazole and CRS3123 as strong candidates for further preclinical development. Future studies should focus on evaluating their pharmacokinetics and efficacy in mammalian models to determine their potential for clinical translation. Given the urgent need for new treatment options against VRE, these compounds warrant further investigation as viable alternatives to current therapies.

## References

1. Guzman Prieto AM, van Schaik W, Rogers MRC, Coque TM, Baquero F, Corander J, et al. Global Emergence and Dissemination of Enterococci as Nosocomial Pathogens: Attack of the Clones? Frontiers in Microbiology. 2016;7.

2. Tripathi A, Shukla SK, Singh A, Prasad KN. Prevalence, outcome and risk factor associated with vancomycin-resistant Enterococcus faecalis and Enterococcus faecium at a Tertiary Care Hospital in Northern India. Indian Journal of Medical Microbiology. 2016;34(1):38–45.

3. Ahmed MO, Baptiste KE. Vancomycin-Resistant Enterococci: A Review of Antimicrobial Resistance Mechanisms and Perspectives of Human and Animal Health. Microbial Drug Resistance. 2017;24(5):590–606.

4. Kaur J, Cao X, Abutaleb NS, Elkashif A, Graboski AL, Krabill AD, et al. Optimization of Acetazolamide-Based Scaffold as Potent Inhibitors of Vancomycin-Resistant Enterococcus. J Med Chem. 2020;63(17):9540–62.

5. An W, Holly KJ, Nocentini A, Imhoff RD, Hewitt CS, Abutaleb NS, et al. Structure-activity relationship studies for inhibitors for vancomycin-resistant Enterococcus and human carbonic anhydrases. J Enzyme Inhib Med Chem. 2022;37(1):1838–44.

6. Tacconelli E, Carrara E, Savoldi A, Harbarth S, Mendelson M, Monnet DL, et al. Discovery, research, and development of new antibiotics: the WHO priority list of antibiotic-resistant bacteria and tuberculosis. The Lancet Infectious Diseases. 2018;18(3):318–27.

7. Abutaleb NS, Shrinidhi A, Bandara AB, Seleem MN, Flaherty DP. Evaluation of 1,3,4-Thiadiazole Carbonic Anhydrase Inhibitors for Gut Decolonization of Vancomycin-Resistant Enterococci. ACS Medicinal Chemistry Letters. 2023;14(4):487–92.

8. Britt NS, Potter EM, Patel N, Steed ME. Effect of Continuous and Sequential Therapy among Veterans Receiving Daptomycin or Linezolid for Vancomycin-Resistant Enterococcus faecium Bacteremia. Antimicrob Agents Chemother. 2017;61(5).

9. Abou Hassan OK, Karnib M, El-Khoury R, Nemer G, Ahdab-Barmada M, BouKhalil P. Linezolid Toxicity and Mitochondrial Susceptibility: A Novel Neurological Complication in a Lebanese Patient. Front Pharmacol. 2016;7:325.

10. Baddour LM, Wilson WR, Bayer AS, Fowler VG, Jr., Tleyjeh IM, Rybak MJ, et al. Infective Endocarditis in Adults: Diagnosis, Antimicrobial Therapy, and Management of Complications: A Scientific Statement for Healthcare Professionals From the American Heart Association. Circulation. 2015;132(15):1435–86.

11. Mohammad H, Younis W, Chen L, Peters CE, Pogliano J, Pogliano K, et al. Phenylthiazole Antibacterial Agents Targeting Cell Wall Synthesis Exhibit Potent Activity in Vitro and in Vivo against Vancomycin-Resistant Enterococci. Journal of Medicinal Chemistry. 2017;60(6):2425–38.

12. Donabedian SM, Perri MB, Vager D, Hershberger E, Malani P, Simjee S, et al. Quinupristin-Dalfopristin Resistance in Enterococcus faecium Isolates from Humans, Farm Animals, and Grocery Store Meat in the United States. Journal of Clinical Microbiology. 2006;44(9):3361–5.

13. Gonzales RD, Schreckenberger PC, Graham MB, Kelkar S, DenBesten K, Quinn JP. Infections due to vancomycin-resistant Enterococcus faecium resistant to linezolid. The Lancet. 2001;357(9263):1179.

14. Munoz-Price LS, Lolans K, Quinn JP. Emergence of Resistance to Daptomycin during Treatment of Vancomycin-Resistant Enterococcus faecalis Infection. Clinical Infectious Diseases. 2005;41(4):565–6.

15. Isaac S, Flor-Duro A, Carruana G, Puchades-Carrasco L, Quirant A, Lopez-Nogueroles M, et al. Microbiome-mediated fructose depletion restricts murine gut colonization by vancomycin-resistant Enterococcus. Nat Commun. 2022;13(1):7718.

16. Abutaleb NS, Seleem MN. Antivirulence activity of auranofin against vancomycin-resistant enterococci: in vitro and in vivo studies. International Journal of Antimicrobial Agents. 2020;55(3):105828.

17. Younis W, AbdelKhalek A, Mayhoub AS, Seleem MN. In Vitro Screening of an FDA-Approved Library Against ESKAPE Pathogens. Curr Pharm Des. 2017;23(14):2147–57.

18. Mohammad H, AbdelKhalek A, Abutaleb NS, Seleem MN. Repurposing niclosamide for intestinal decolonization of vancomycin-resistant enterococci. International Journal of Antimicrobial Agents. 2018;51(6):897–904.

19. Brown D. Antibiotic resistance breakers: can repurposed drugs fill the antibiotic discovery void? Nature Reviews Drug Discovery. 2015;14(12):821–32.

20. Abutaleb NS, Elhassanny AEM, Flaherty DP, Seleem MN. In vitro and in vivo activities of the carbonic anhydrase inhibitor, dorzolamide, against vancomycin-resistant enterococci. PeerJ. 2021;9:e11059.

21. Mohamed MF, Brezden A, Mohammad H, Chmielewski J, Seleem MN. Targeting biofilms and persisters of ESKAPE pathogens with P14KanS, a kanamycin peptide conjugate. Biochimica et Biophysica Acta - General Subjects. 2017;1861(4):848–59.

22. Adan A, Kiraz Y, Baran Y. Cell Proliferation and Cytotoxicity Assays. Curr Pharm Biotechnol. 2016;17(14):1213–21.

23. Mohamed MF, Brezden A, Mohammad H, Chmielewski J, Seleem MN. A short D-enantiomeric antimicrobial peptide with potent immunomodulatory and antibiofilm activity against multidrug-resistant Pseudomonas aeruginosa and Acinetobacter baumannii. Scientific Reports. 2017;7(1):6953.

24. Mohamed MF, Hamed MI, Panitch A, Seleem MN. Targeting Methicillin-Resistant Staphylococcus aureus with Short Salt-Resistant Synthetic Peptides. Antimicrobial Agents and Chemotherapy. 2014;58(7):4113–22.

25. Moy TI, Ball AR, Anklesaria Z, Casadei G, Lewis K, Ausubel FM. Identification of novel antimicrobials using a live-animal infection model. Proceedings of the National Academy of Sciences. 2006;103(27):10414–9.

26. Tampakakis E, Okoli I, Mylonakis E. A C. elegans-based, whole animal, in vivo screen for the identification of antifungal compounds. Nature Protocols. 2008;3(12):1925–31.

27. Kim Y, Mylonakis E. Caenorhabditis elegans immune conditioning with the probiotic bacterium Lactobacillus acidophilus strain NCFM enhances gram-positive immune responses. Infect Immun. 2012;80(7):2500–8.

28. Garsin DA, Sifri CD, Mylonakis E, Qin X, Singh KV, Murray BE, et al. A simple model host for identifying Gram-positive virulence factors. Proc Natl Acad Sci U S A. 2001;98(19):10892–7.

29. Collins DA, Riley TV. Ridinilazole: a novel, narrow-spectrum antimicrobial agent targeting Clostridium (Clostridioides) difficile. Letters in Applied Microbiology. 2022;75(3):526–36.

30. Critchley IA, Green LS, Young CL, Bullard JM, Evans RJ, Price M, et al. Spectrum of activity and mode of action of REP3123, a new antibiotic to treat Clostridium difficile infections. J Antimicrob Chemother. 2009;63(5):954–63.

31. Ochsner UA, Young CL, Stone KC, Dean FB, Janjic N, Critchley IA. Mode of action and biochemical characterization of REP8839, a novel inhibitor of methionyl-tRNA synthetase. Antimicrob Agents Chemother. 2005;49(10):4253–62.

32. Collins DA, Riley TV. Ridinilazole: a novel, narrow-spectrum antimicrobial agent targeting Clostridium (Clostridioides) difficile. Lett Appl Microbiol. 2022;75(3):526–36.

33. Cho JC, Crotty MP, Pardo J. Ridinilazole: a novel antimicrobial for Clostridium difficile infection. Ann Gastroenterol. 2019;32(2):134–40.

34. Nayak SU, Griffiss JM, Blumer J, O’Riordan MA, Gray W, McKenzie R, et al. Safety, Tolerability, Systemic Exposure, and Metabolism of CRS3123, a Methionyl-tRNA Synthetase Inhibitor Developed for Treatment of Clostridium difficile, in a Phase 1 Study. Antimicrobial Agents and Chemotherapy. 2017;61(8):10.1128/aac.02760-16.

35. Mason CS, Avis T, Hu C, Nagalingam N, Mudaliar M, Coward C, et al. The Novel DNA Binding Mechanism of Ridinilazole, a Precision *Clostridiodes difficile* Antibiotic. Antimicrobial Agents and Chemotherapy. 2023;67(5):e01563–22.

36. Lomeli BK, Galbraith H, Schettler J, Saviolakis GA, El-Amin W, Osborn B, et al. Multiple-Ascending-Dose Phase 1 Clinical Study of the Safety, Tolerability, and Pharmacokinetics of CRS3123, a Narrow-Spectrum Agent with Minimal Disruption of Normal Gut Microbiota. Antimicrob Agents Chemother. 2019;64(1).

